# Brazilian bryophytes: a national-scale comprehensive dataset of species richness and geographic distribution

**DOI:** 10.64898/2026.11.01.763487

**Authors:** Jailton Venilson Ferreira da Silva, Jhonyd Jhonata de Oliveira Marmo, Kleberson Jonata da Cruz Tavares, Hermeson Cassiano de Oliveira, Mércia Patrícia Pereira Silva

## Abstract

Bryophytes (Anthocerotophyta, Marchantiophyta, and Bryophyta) remain under-sampled in large-scale biodiversity databases in Brazil, despite their ecological importance. In Brazil, Flora e Funga do Brasil recognizes 1,618 bryophyte species distributed across six biomes that are increasingly threatened by anthropogenic pressures such as deforestation, agricultural expansion, and climate change. Nevertheless, knowledge of species geographic distributions remains fragmented and based on scattered records. Flora e Funga do Brasil serves as the official taxonomic reference for the country, providing the accepted species list and serving as a baseline for biodiversity studies. However, as it was designed as a nomenclatural and taxonomic checklist, it does not integrate primary occurrence records with geographic coordinates and collection dates. Consequently, occurrence data remain dispersed across herbaria, digital repositories, and the scientific literature, without dynamic updates linked to the official checklist. This disconnect between the taxonomic framework and primary occurrence records limits macroecological and biogeographical analyses at national scales, which depend on standardized georeferenced biodiversity databases. To address this gap, we compiled 102,528 bryophyte occurrence records from herbarium specimens, using data from GBIF and the speciesLink network, complemented by field collections from the Caatinga and records from the Cryptogamic Herbarium of the State University of Piauí. All records were georeferenced and curated, including verification of inconsistent coordinates, removal of duplicates, and taxonomic standardization using Bryonames. When available, information such as collection date and collector was retained. The dataset comprises 1,820 species, 125 families, and 442 genera. Bryophyta (mosses) was the most diverse group (1,056 spp., 84 families, 310 genera), followed by Marchantiophyta (745 spp., 37 families, 124 genera) and Anthocerotophyta (19 spp., four families, seven genera). Among mosses, the most species-rich families were Sphagnaceae (79 spp.), Fissidentaceae (76 spp.), and Pottiaceae (76 spp.). Among liverworts, Lejeuneaceae dominated with 335 species, followed by Lepidoziaceae (58 spp.) and Plagiochilaceae (51 spp.). Among hornworts, Anthocerotaceae, Dendrocerotaceae, and Notothyladaceae each included six species. Regionally, the Southeast showed the highest richness (1,338 spp.), followed by the South (974 spp.), Central-West (937 spp.), Northeast (897 spp.), and North (763 spp.). Regarding biomes, the Atlantic Forest (1,523 spp.) and Amazon (1,054 spp.) were the most species-rich, followed by the Cerrado (810 spp.), Caatinga (671 spp.), Pampa (269 spp.), and Pantanal (61 spp.). These patterns reflect both ecological gradients and historical sampling biases. This dataset bridges the gap between the official taxonomic checklist and primary occurrence data by integrating multiple sources into a standardized, openly accessible database. It provides a robust tool for studies in macroecology, biogeography, and conservation, enabling analyses of spatial patterns of species richness, endemism, and taxonomic, functional, and phylogenetic diversity across environmental gradients. It also supports species distribution modeling, climate change impact assessments, biodiversity hotspot identification, and conservation planning at regional and national scales. Users are encouraged to cite this article when using the dataset and to report downstream applications derived from it.

Open Research statement: The data is provided as a private submission for peer review via the following link:

## Introduction

Studies on biodiversity in tropical plant communities have predominantly focused on trees (Condit et al., 2002; Eisenlohr et al., 2013; Gentry, 1988; Oliveira & Daly, 1999; Oliveira-Filho et al., 2005; Pitman et al., 2001, 2002; Ter Steege et al., 2003), whereas bryophytes (Anthocerotophyta—hornworts, Marchantiophyta—liverworts, and Bryophyta—mosses) have often been overlooked despite their ecological importance. These plants contribute to water retention, facilitate the establishment of vascular plants (Welch, 1948), and provide essential microhabitats for a wide range of organisms (Gradstein et al., 2001; Glime, 2007).

In Brazil, 1,618 bryophyte species are currently recognized (18 hornworts, 702 liverworts, and 898 mosses; Flora e Funga do Brasil, 2025), distributed across different biomes that are increasingly threatened by anthropogenic pressures such as deforestation, climate change, agricultural expansion, wildfires, and urbanization (Hirota, 2003; INPE, 2016; Imazon, 2020; Walter & Sevilha, 2019). Despite these threats, the country’s unique combination of humid and dry forests still provides suitable conditions for bryophyte development. However, knowledge of their distribution remains fragmented and incomplete (Aires & Bordin, 2024; Carvalho et al., 2025; Lima & Peralta, 2021; Marmo & Silva, 2025; Silva et al., 2024, 2025; Sousa et al., 2024).

Despite significant advances in the knowledge of the Brazilian bryoflora over recent decades, information on species occurrences remains scattered across biological collections, digital herbaria, regional inventories, and scientific publications. In this context, Flora e Funga do Brasil (2025) represents the first official taxonomic framework for the country, providing a comprehensive checklist of recognized species (Forzza et al., 2012). However, primary occurrence records including geographic coordinates, collector information, collection dates, and substrate data remain dispersed among biological collections, digital herbaria, and the specialized literature. Initiatives such as SpeciesLink and GBIF have integrated part of these records (Canhos et al., 2004; GBIF.org, 2025), but they do not replace a consolidated and standardized national-scale database. This limitation stems from the fact that Flora e Funga do Brasil was not designed as a dynamic repository of occurrence records, while taxonomic updates and herbarium data continue to follow independent update cycles.

Furthermore, collecting effort has historically been uneven across Brazilian regions and biomes, generating spatial and taxonomic biases that hinder a comprehensive understanding of species distribution and richness patterns (Costa & Peralta, 2015). Consequently, the establishment of large, standardized databases is crucial for advancing studies in macroecology, biogeography, conservation, and environmental change monitoring (Antunes et al., 2022; Costa-Pinto et al., 2022; Costa et al., 2020; Hasui et al., 2017; Ramos et al., 2019; Travers & Dunnell, 2009).

In this study, we present a dataset comprising more than 100,000 bryophyte occurrence records from Brazil, encompassing all three bryophyte phyla across a wide range of biomes. The dataset integrates taxonomic, spatial, and temporal information and provides a valuable resource for analyses of species richness, geographic distribution, taxonomic representativeness, and collecting effort over time. Furthermore, it serves as a foundation for future research in biodiversity, conservation, macroecology, biogeography, and ecological modeling.

## Description

A total of 102,528 bryophyte occurrence records were compiled for Brazil, encompassing representatives of all three phyla: Anthocerotophyta, Bryophyta, and Marchantiophyta. The records show a broad distribution across the Brazilian territory, with the highest concentration of occurrences in the Atlantic Forest and along coastal regions of the Southeast and South (Figure 1). In contrast, lower record densities were observed in the Pantanal and Pampa biomes, as well as in inland areas of the Northern region.

**Figure 1.**
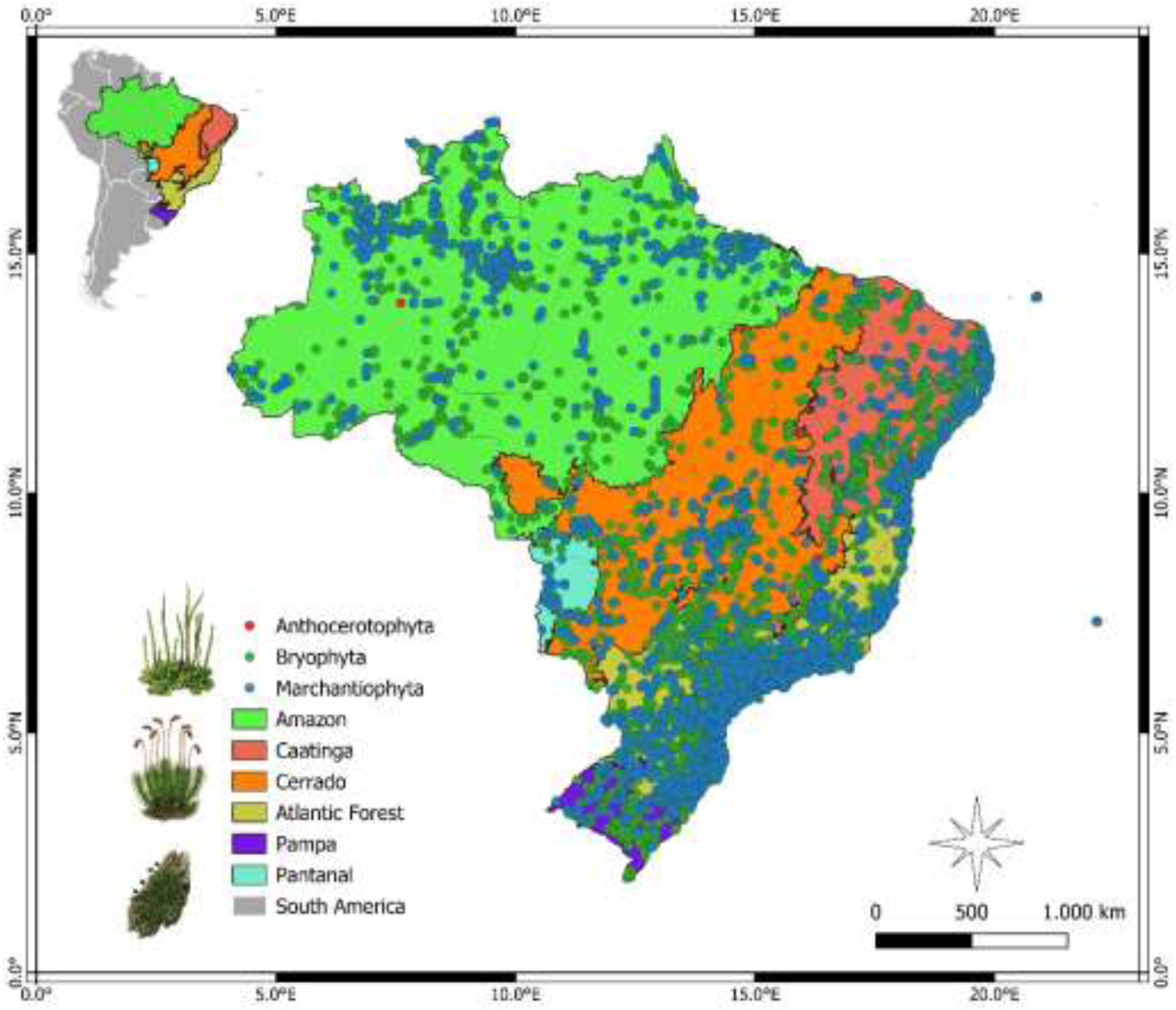
Distribution of bryophyte occurrence records (Anthocerotophyta, Bryophyta, and Marchantiophyta) across Brazilian biomes.

To better understand spatial distribution patterns, occurrence records were subsequently analyzed separately for each bryophyte phylum. Bryophyta accounted for 68,478 records across Brazil (Figure 2), whereas Marchantiophyta comprised 33,594 records (Figure 3). Bryophyta and Marchantiophyta exhibited the broadest geographic distributions and the highest numbers of records throughout the country. In contrast, Anthocerotophyta was less well represented, with only 455 records and a more restricted spatial distribution (Figure 4).

**Figure 2.**
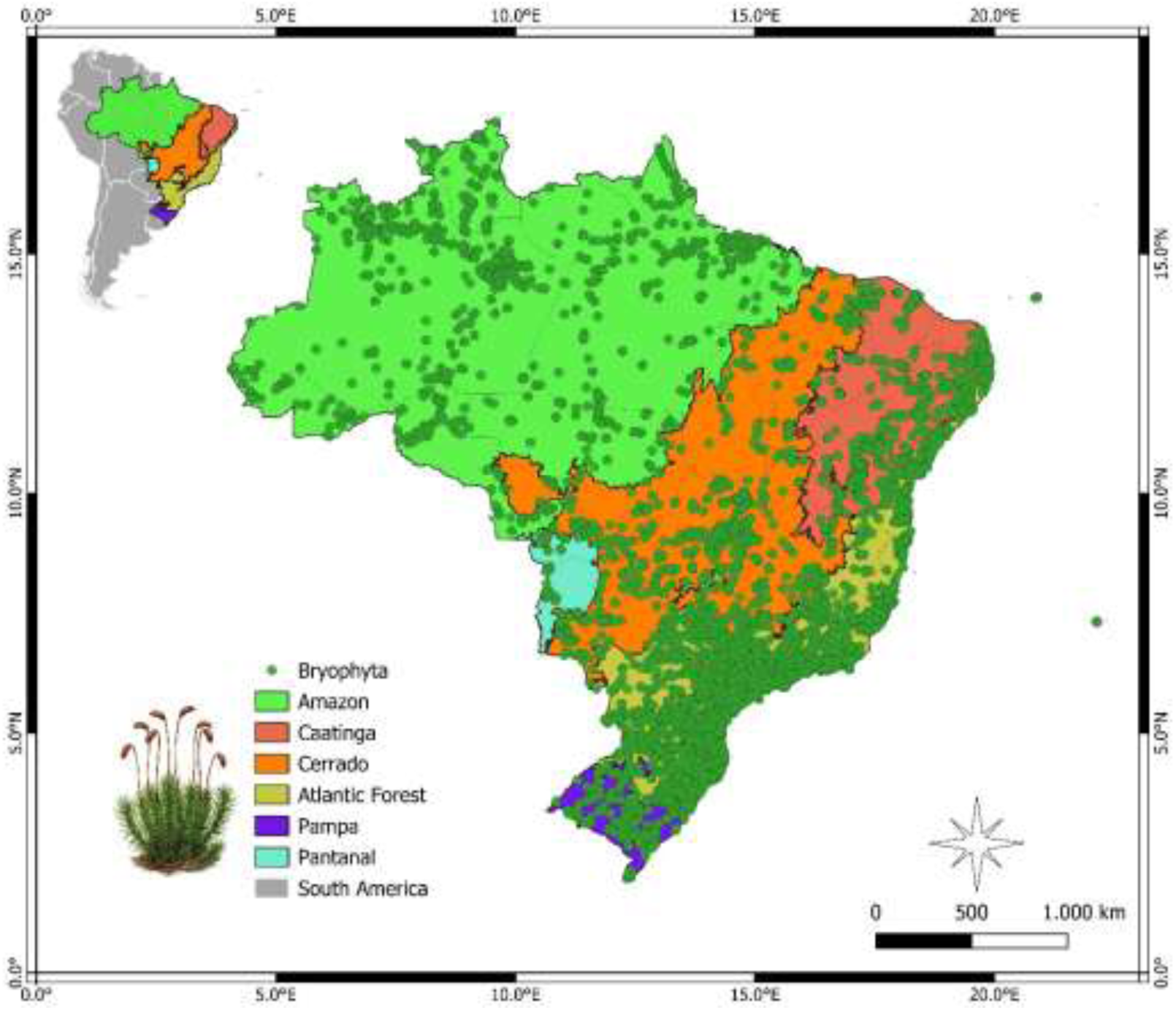
Spatial distribution of moss (Bryophyta) occurrence records across the Brazilian biomes.

**Figure 3.**
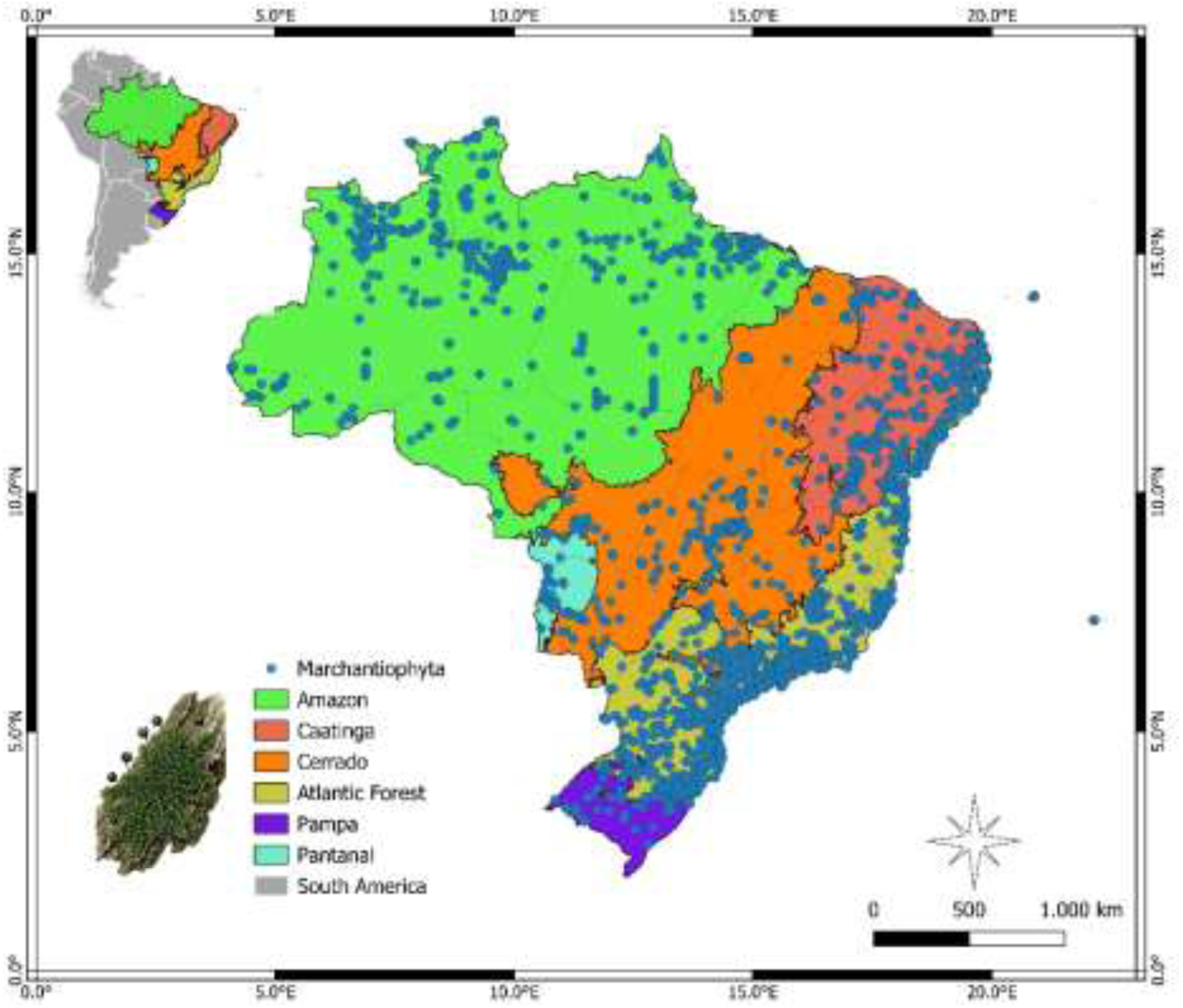
Spatial distribution of liverwort (Marchantiophyta) occurrence records across the Brazilian biomes.

**Figure 4.**
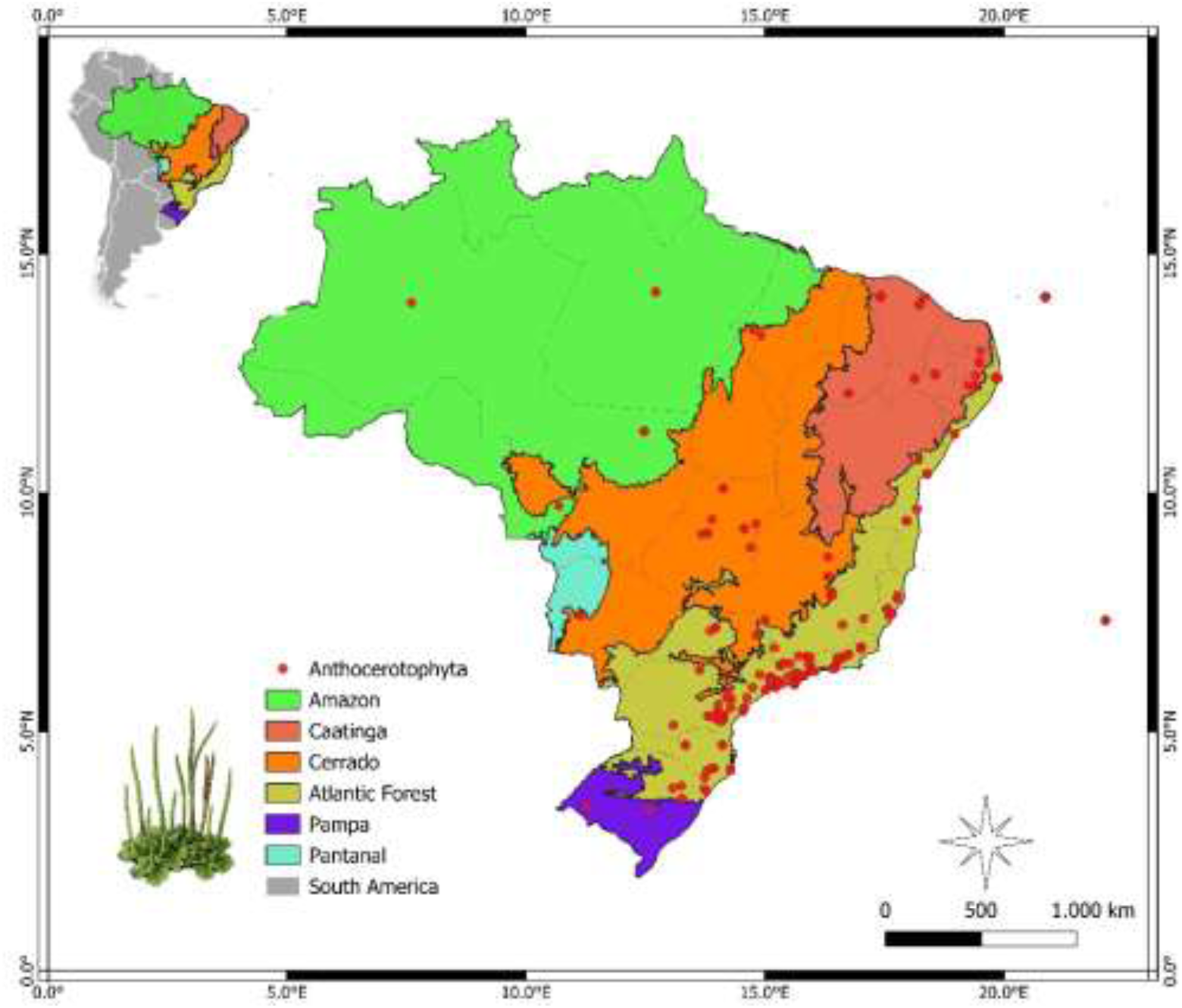
Spatial distribution of hornwort (Anthocerotophyta) occurrence records across the Brazilian biomes.

The distribution of records was also analyzed across the different Brazilian biomes for all three bryophyte phyla. The Atlantic Forest exhibited the highest numbers of occurrence records and species richness, followed by the Amazon, Caatinga, and Cerrado. In contrast, the Pantanal and Pampa showed the lowest representation in terms of both record numbers and species richness. Bryophyta and Marchantiophyta were widely distributed across all biomes, particularly in humid forested regions, whereas Anthocerotophyta displayed a more restricted distribution concentrated in specific areas.

The complete dataset comprises 1,820 identified species distributed across 125 families and 442 genera. Bryophyta was the most diverse division, with 1,056 species, 310 genera, and 84 families, followed by Marchantiophyta, with 745 species, 124 genera, and 37 families, and Anthocerotophyta, with 19 species, seven genera, and four families. Among the mosses (Bryophyta), the most species-rich families were Sphagnaceae (79 spp.), Fissidentaceae (76 spp.), and Pottiaceae (76 spp.). Among the liverworts (Marchantiophyta), Lejeuneaceae was the dominant family, comprising 335 species, followed by Lepidoziaceae (58 spp.) and Plagiochilaceae (51 spp.). Among the hornworts (Anthocerotophyta), the most species-rich families were Anthocerotaceae, Dendrocerotaceae, and Notothyladaceae, each represented by six species.

The Atlantic Forest contained the highest number of bryophyte records, totaling 54,237 occurrences, followed by the Amazon, with 18,560 records. The Caatinga and Cerrado were also well represented, accounting for 14,561 and 13,856 records, respectively. The lowest numbers of records were observed in the Pampa, with 954 occurrences, and the Pantanal, with only 146 records. Species richness was likewise highest in the Atlantic Forest, where 1,523 species were recorded, followed by the Amazon with 1,054 species. The Cerrado harbored 810 species, whereas the Caatinga contained 671 recorded species. The lowest species richness values were found in the Pampa, with 269 species, and the Pantanal, with only 61 recorded species.

The Southeast region contained the highest number of bryophyte records, totaling 32,069 occurrences. It was followed by the Northeast, with 30,160 records. The North accounted for 15,288 records, whereas the South and Central-West regions contributed 12,607 and 12,403 records, respectively. Species richness was also highest in the Southeast, with 1,338 recorded species, followed by the South, with 974 species. The Central-West harbored 937 species, while the Northeast accounted for 897 species. The lowest species richness was observed in the North, with 763 recorded species.

At the state level, Bahia exhibited the highest number of bryophyte records, totaling 18,367 occurrences, followed by São Paulo, with 15,292 records. Amazonas accounted for 7,598 records, while Minas Gerais and Rio de Janeiro contributed 6,741 and 5,291 records, respectively. Other states with substantial representation included Pará (4,785 records), Espírito Santo (4,745), Mato Grosso (4,728), Rio Grande do Sul (4,376), Paraná (4,305), and Santa Catarina (3,926). Intermediate numbers of records were observed in Goiás (3,433 records), Pernambuco (3,347), the Federal District (3,312), Ceará (2,809), and Piauí (2,596). The lowest numbers of records were found in Rondônia (1,126), Alagoas (1,072), Paraíba (932), Mato Grosso do Sul (930), Roraima (769), Sergipe (671), Acre (558), Amapá (380), Rio Grande do Norte (192), Maranhão (174), and Tocantins (72 records).

The highest species richness was recorded in São Paulo, with 999 species, followed by Minas Gerais (826 species), Rio de Janeiro (810), Bahia (770), and Mato Grosso (736). High species richness was also observed in Paraná (667 species), Santa Catarina (618), Rio Grande do Sul (610), Amazonas (591), and Espírito Santo (528). Intermediate richness values were recorded in Pará (410 species), Pernambuco (398), Goiás (382), the Federal District (332), Ceará (285), Roraima (235), and Alagoas (221). The lowest richness values were observed in Paraíba (194 species), Piauí (187), Rondônia (184), Mato Grosso do Sul (180), Acre (145), Sergipe (125), Amapá (90), Maranhão (77), Tocantins (48), and Rio Grande do Norte, with only 41 recorded species.

We analyzed bryophyte records across five temporal periods and observed a marked increase in the number of records over time. Between 1800–1850 and 1901–1950, the number of records remained relatively low, indicating limited sampling effort during the early stages of bryological exploration in Brazil. From 1951–2000 onward, there was a sharp increase in recorded occurrences, while the 2001–2025 period exhibited the highest values, exceeding 45,000 records (Figure 5). This pattern likely reflects the expansion of floristic surveys, the growth in the number of bryology specialists, the strengthening of biological collections, and the increasing digitization of Brazilian herbaria.

**Figure 5.**
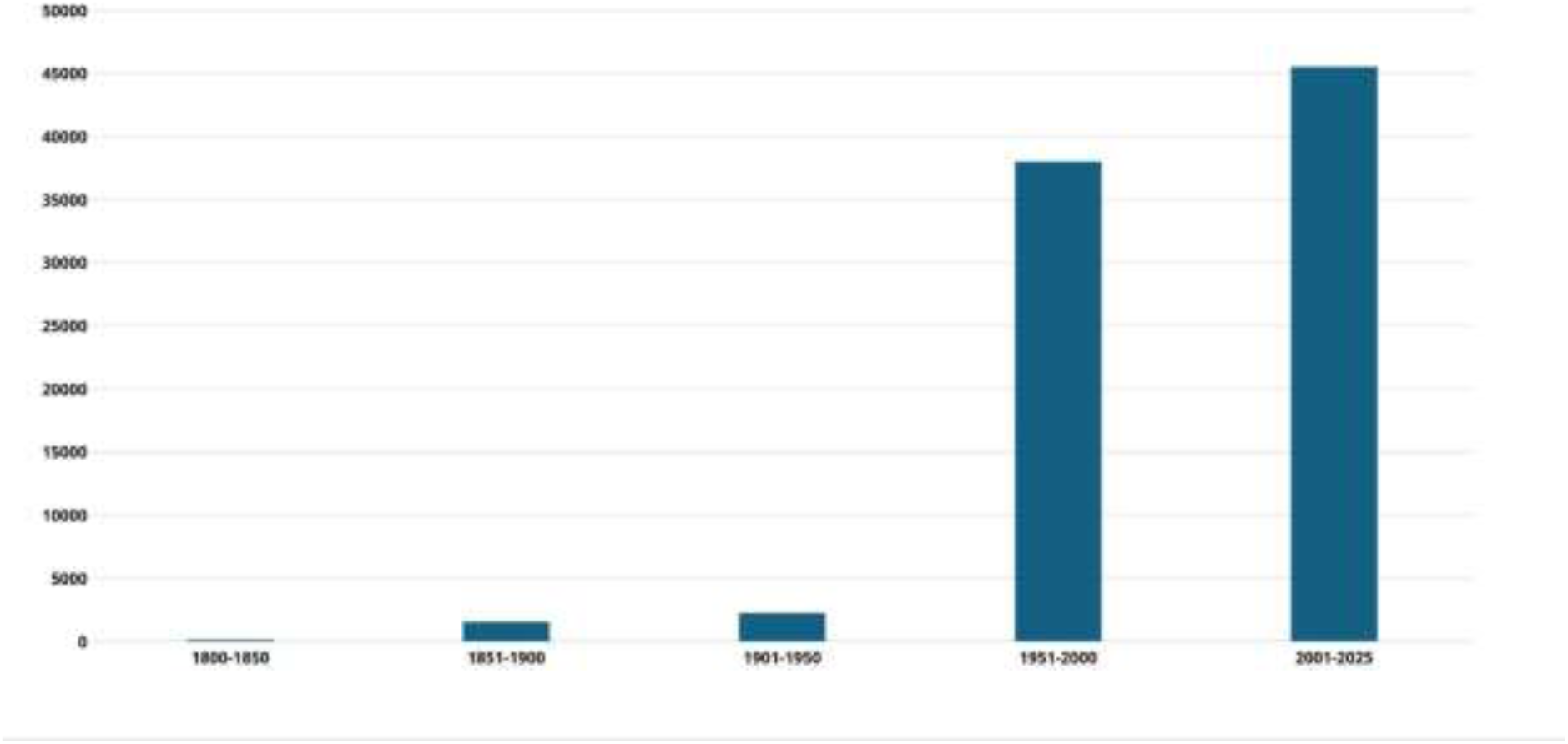
Temporal distribution of bryophyte occurrence records in Brazil from 1800 to 2025.

We also analyzed bryophyte species richness over time. The results show a progressive increase in the number of species recorded in Brazil throughout the studied periods. Between 1800–1850, only a few species had been documented, whereas subsequent periods exhibited a continuous increase in known species richness. The most pronounced increase occurred between 1951–2000, when the number of recorded species exceeded 1,400. During the 2001–2025 period, the upward trend in species richness was maintained, with a slight increase of two species compared to the previous interval. This pattern may be related to differences in sampling effort, temporal variation in data availability, or the absence of recently collected records that have not yet been fully digitized (Figure 6).

**Figure 6.**
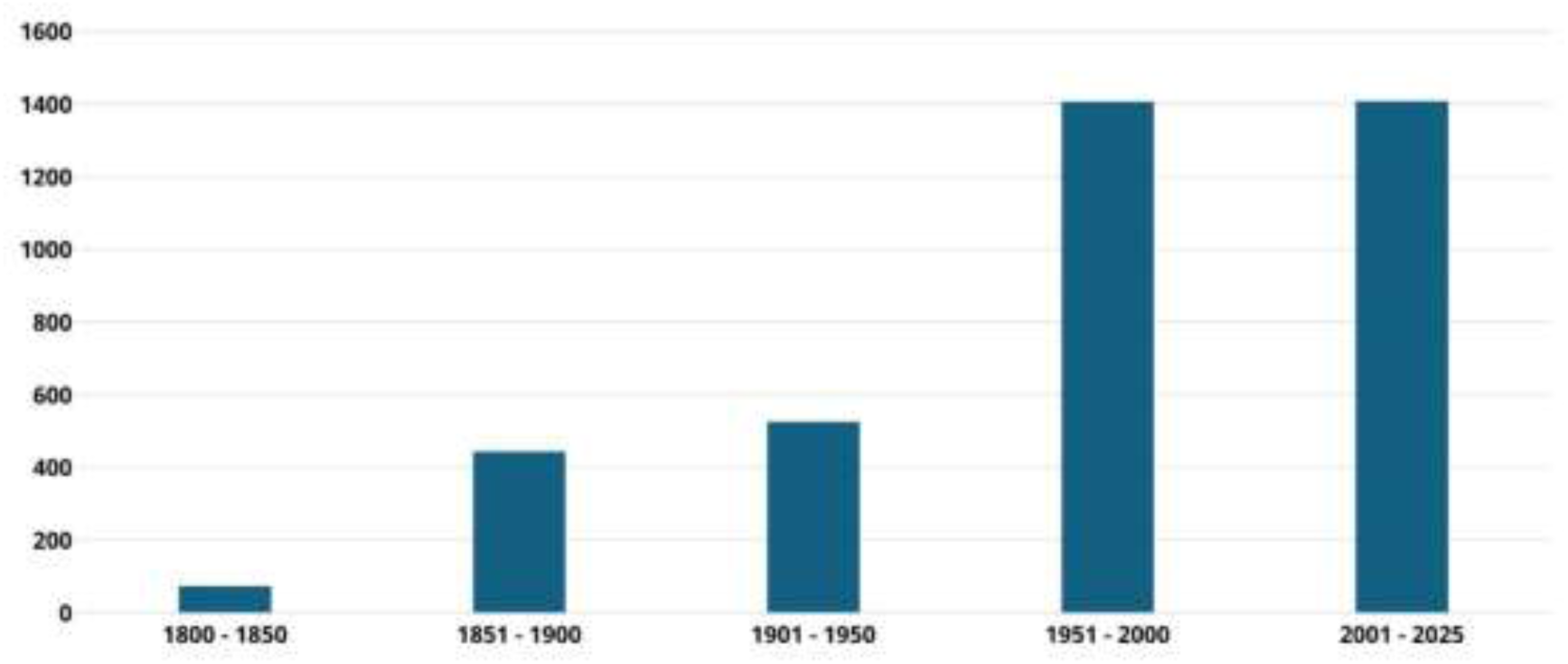
Temporal variation in bryophyte species richness in Brazil from 1800 to 2025.

The number of bryophyte records increased markedly over time across all Brazilian biomes. In the earliest periods (1800–1850 and 1851–1900), records were relatively scarce, with the Atlantic Forest already showing the highest number of documented occurrences. From 1951–2000 onward, a pronounced increase in records was observed across all biomes, becoming even more evident in the 2001–2025 period. The Atlantic Forest consistently concentrated the highest number of records across all recent periods, reaching 27,157 records between 2001–2025, followed by the Cerrado (8,332 records), Caatinga (6,642 records), and Amazon (4,695 records). The Pantanal and Pampa exhibited the lowest numbers of records throughout the entire time series, indicating lower sampling intensity or limited data availability in these biomes (Figure 7).

**Figure 7.**
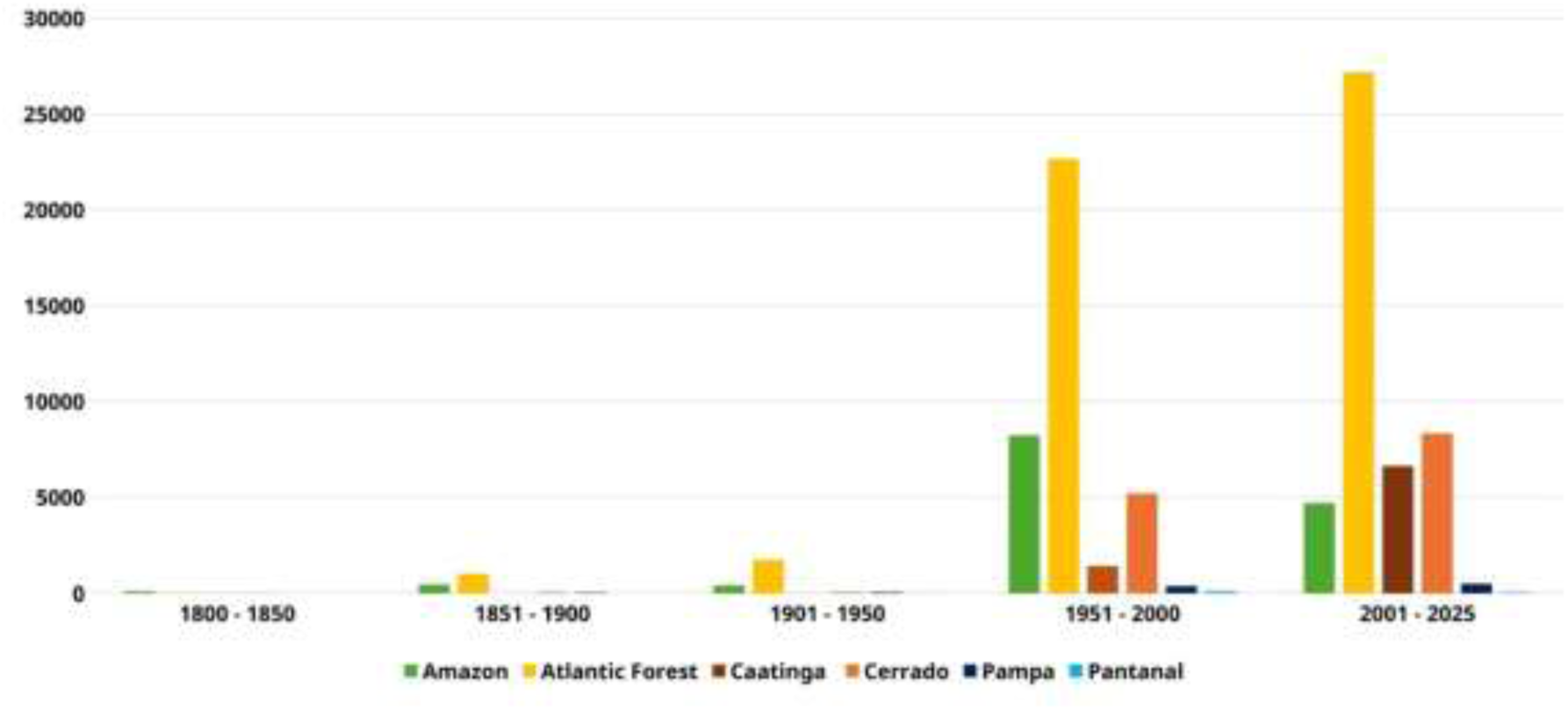
Temporal distribution of bryophyte occurrence records across Brazilian biomes from 1800 to 2025.

Species richness also increased over the analyzed periods, following the growth in the number of records. The Atlantic Forest exhibited the highest richness across all periods, reaching 1,166 species between 2001–2025. It was followed by the Cerrado (557 species), Amazon (522 species), and Caatinga (520 species). Although species richness increased substantially up to the 1951–2000 period, some biomes showed a slight reduction in the number of recorded species between 2001–2025, including the Atlantic Forest (from 1,196 to 1,166 species), the Amazon (from 540 to 522 species), and the Cerrado (from 591 to 557 species). This reduction may reflect differences in sampling effort, taxonomic updates, or data standardization procedures. The Pampa and Pantanal consistently exhibited the lowest richness values throughout the time series, reaching 147 and 21 species, respectively, in the most recent period. In contrast, the Caatinga showed one of the largest relative increases in richness, rising from only two recorded species in 1800–1850 to 520 species in 2001–2025 (Figure 8).

**Figure 8.**
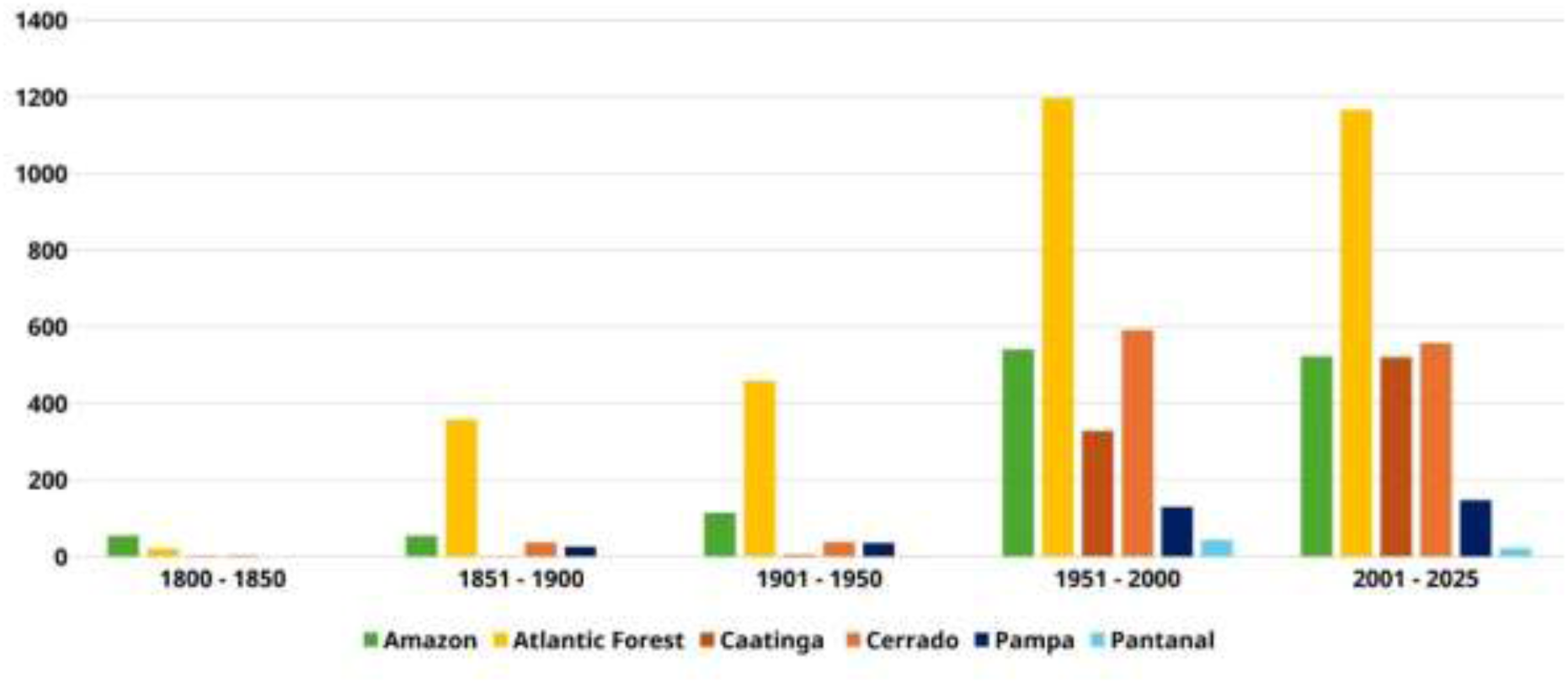
Temporal variation in bryophyte species richness across Brazilian biomes from 1800 to 2025.

The fifteen most representative moss families and genera in the dataset were selected based on both the number of occurrence records and species richness. Sematophyllaceae had the highest number of records (6,938) and 57 species, followed by Leucobryaceae (6,800 records; 66 species), Calymperaceae (5,354; 52 species), and Fissidentaceae (4,634; 76 species). Other well-represented families included Orthotrichaceae, Pylaisiadelphaceae, and Sphagnaceae (Figure 9). At the genus level, Fissidens was the most representative, with 4,634 records and 76 species, followed by Campylopus (4,255 records; 48 species), Syrrhopodon (3,116; 33 species), and Sphagnum (3,100 records; 79 species). Other relevant genera included Octoblepharum, Microcalpe, and Calymperes (Figure 10).

**Figure 9.**
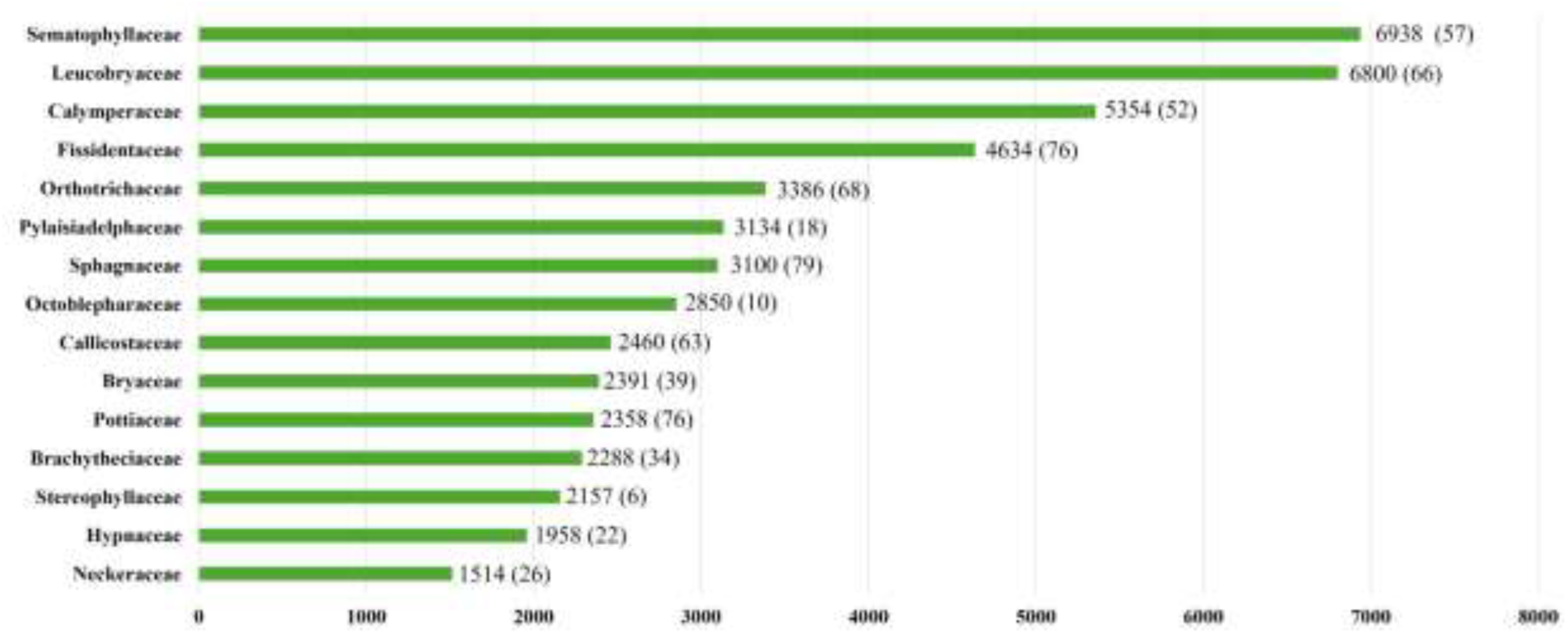
Number of occurrence records for the 15 most representative moss families, with species richness indicated in parentheses.

**Figure 10.**
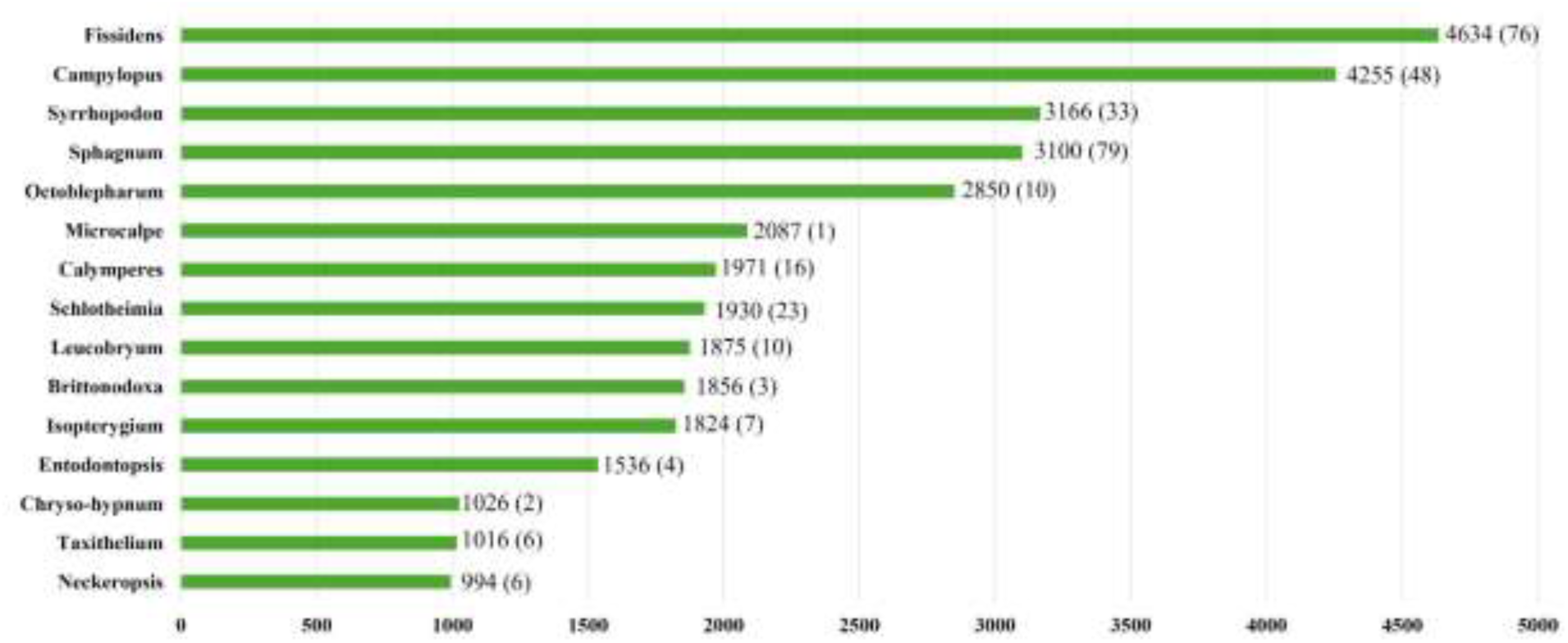
Number of occurrence records for the 15 most representative moss genera, with species richness indicated in parentheses.

As with mosses, the fifteen most representative liverwort families and genera in the dataset were selected based on the number of occurrence records and species richness. Among families, Lejeuneaceae stood out prominently, with 17,648 records and 335 species, followed by Frullaniaceae (3,520 records; 44 species), Plagiochilaceae (2,787; 51 species), and Lepidoziaceae (2,069; 58 species). Other relevant families included Radulaceae, Metzgeriaceae, and Ricciaceae (Figure 11). At the genus level, Lejeunea had the highest number of records, totaling 3,876 occurrences and 68 species, followed by Frullania (3,521 records; 44 species), Cheilolejeunea (3,238; 41 species), and Plagiochila (2,764; 47 species). Additional notable genera included Ceratolejeunea, Radula, Metzgeria, and Cololejeunea (Figure 12).

**Figure 11.**
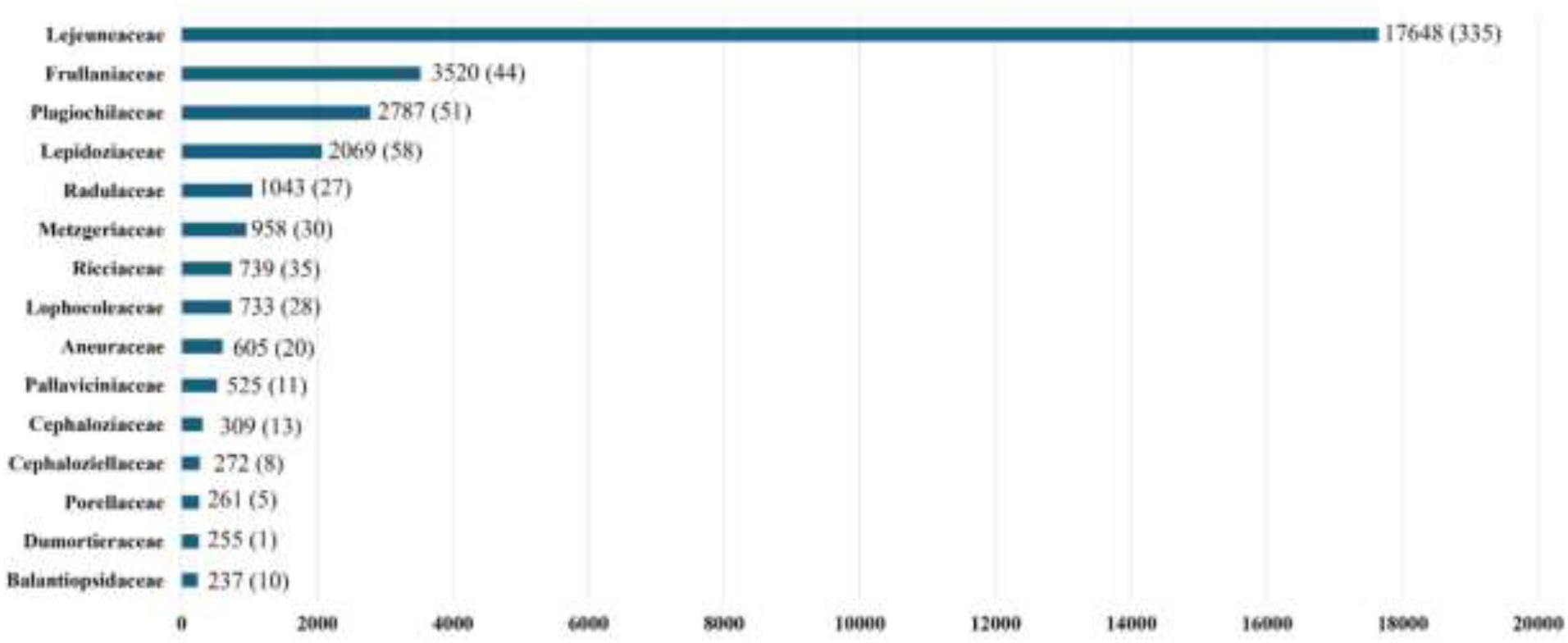
Number of occurrence records for the 15 most representative liverwort families, with species richness indicated in parentheses.

**Figure 12.**
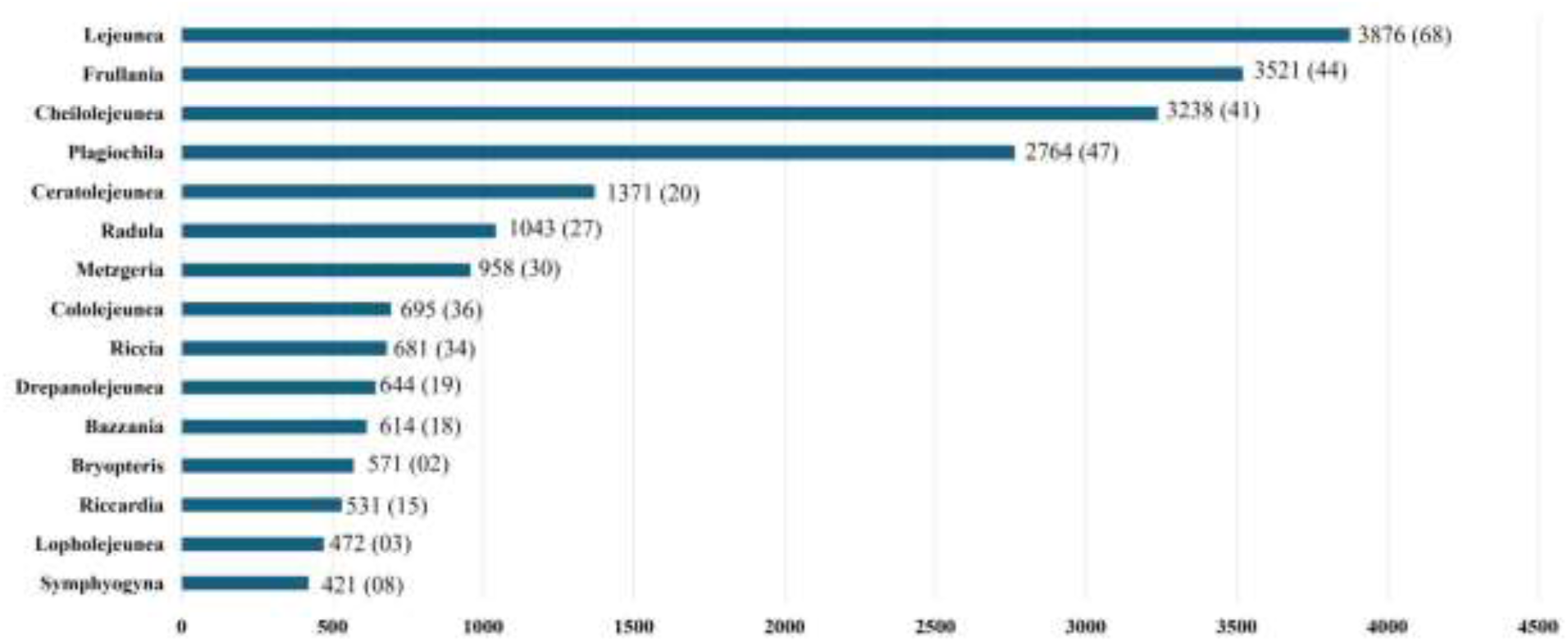
Number of occurrence records for the 15 most representative liverwort genera, with species richness indicated in parentheses.

## Class II. Research origin descriptors

### A. Overall project description

1. Identity: A compilation of bryophyte occurrence data from Brazil.
2. Originators: The Brazilian Bryophytes Project was coordinated by Jailton Venilson Ferreira da Silva (Federal University of Pernambuco, UFPE). All co-authors contributed to the compilation, validation, and organization of the dataset. The research team is primarily composed of investigators from UFPE, with additional collaboration from a researcher at the State University of Piauí (UESPI).
3. Period of study: The dataset covers the period from 1804 to 2025.
4. Objectives: Our objectives were to: (1) compile a comprehensive dataset of Brazilian bryophytes, encompassing the three major groups, hornworts (Anthocerotophyta), liverworts (Marchantiophyta), and mosses (Bryophyta), based on occurrence records derived from field collections and herbarium specimens; and (2) characterize the distribution of Brazilian bryophytes in order to identify priorities for future sampling efforts. To our knowledge, this study represents the first nationwide bryophyte dataset for Brazil integrating occurrence records with information on biomes and geographic regions.
5. 5. Abstract: Same as above.
6. Sources of funding: The compilation of this dataset was supported by Coordenação de Aperfeiçoamento de Pessoal de Nível Superior (CAPES; Finance Code 001) and Conselho Nacional de Desenvolvimento Científico e Tecnológico (CNPq; Universal Grant No. 409503/2025-8 and PPBio-RABECA Network Grant No. 441209/2023-8).

### B. Specific subproject description

#### 1. Site description

Brazil is recognized as a country with extensive forest cover, comprising approximately 500 million hectares of natural and planted forests, which correspond to about 59% of its national territory. This places Brazil as the country with the second-largest forest area in the world, surpassed only by Russia. In addition, Brazil covers approximately 8.5 million km², extending between latitudes 5°16’N and 33°45’S and longitudes 34°47’W and 73°59’W (IBGE, 2025; Serviço Florestal Brasileiro, 2019). Brazil’s natural resources are under considerable pressure from large-scale agricultural expansion, mining activities, and timber extraction, creating major challenges for balancing economic development with environmental conservation and the sustainable use of ecosystems (Laurance et al., 2001). The country encompasses six major biomes, Amazon, Cerrado, Caatinga, Atlantic Forest, Pantanal, and Pampa, two of which are recognized as global biodiversity hotspots: the Cerrado and the Atlantic Forest. Together, these biomes harbor approximately 2% of the world’s endemic plant and vertebrate species and account for more than two-thirds of the species richness recorded in Brazil (Myers et al., 2000). These biomes are characterized by distinct vegetation formations, climatic conditions, and precipitation regimes. The Atlantic Forest and Amazon are predominantly humid forest ecosystems, whereas the Cerrado and Caatinga form a broad corridor of seasonally dry vegetation. The Pampa, located in southern Brazil, is dominated by grassland formations, while the Pantanal is characterized by extensive floodplain vegetation and seasonal inundation dynamics (Oliveira-Filho & Fontes, 2000; Oliveira-Filho & Ratter, 2002; Oliveira-Filho et al., 2005; Silva et al., 2017; Serviço Florestal Brasileiro, 2019). Brazilian biomes have been subjected to intense anthropogenic pressures, particularly as a result of deforestation, climate change, agricultural expansion, wildfires, and urban growth. These impacts have contributed to biodiversity loss, habitat fragmentation, and alterations in natural ecological processes (Hirota, 2003; Imazon, 2020; INPE, 2016; Walter & Sevilha, 2019). In this context, comprehensive inventories and biodiversity databases are essential for understanding patterns of biodiversity distribution in Brazil, particularly for still under-sampled groups such as bryophytes.

#### 2. Experimental or sampling design

Data compilation: All data were obtained from records of specimens deposited in herbaria, using the Global Biodiversity Information Facility (GBIF; GBIF.org [2025a, 2025b, 2025c]) as the primary data source due to its broad international coverage. Additionally, the speciesLink network was consulted to incorporate data from Brazilian collections not indexed in GBIF. The dataset was further complemented with field collections conducted in the Caatinga domain, as well as records from the Cryptogam Herbarium of the State University of Piauí (UESPI), which is not currently indexed in any online biodiversity platform.

#### 3. Research methods

The database of moss, liverwort, and anthocerotophyte occurrences was compiled based on records of specimens deposited in herbaria, using the Global Biodiversity Information Facility (GBIF; GBIF.org [2025a, 2025b, 2025c]) as the primary source, selected for its broad international coverage. In addition, the speciesLink network (splink.org.br) was consulted to integrate data from national collections not indexed in GBIF. The search and data extraction followed standardized criteria. In GBIF, the search was limited to the terms “Bryophyta,” “Marchantiophyta,” and “Anthocerotophyta.” The search was conducted individually for each division, filtering for specimens preserved in scientific collections (Record Bases field) with available geographic coordinates (Location field). For speciesLink, the filters were set to “Preserved specimen” in the record base. The terms referring to bryophytes in the “Phylum” section were: “Bryophytes,” “Bryophyta,” “Marchantiophyta,” “Anthocerotophyta,” “Moss,” “Hepatics,” “Bryidae,” “Hepatophyta,” and “Marchantiidae.” To avoid pseudo-duplication of data across platforms, a manual review of the records was conducted, excluding from the speciesLink dataset those specimens already obtained via GBIF. After consolidation, a taxonomic quality filter was applied: only samples identified or collected by bryophyte specialists were retained in the dataset. This procedure aims to mitigate identification errors, ensuring the taxonomic robustness essential for studies of distribution patterns and ecological modeling. Subsequently, the geographic coordinates of each record included in the dataset were individually verified using Google Maps to confirm they matched the location provided in the records. The nomenclatural updating of species was based on the BryoNames database (bryonames.org/) using the nameMatch function from the U.Taxonstand package in R Studio v4.5 (R Core Team 2025). When the verified species had multiple synonyms, the Flora and Funga of Brazil database (floradobrasil.jbrj.gov.br) was consulted to determine which of these synonyms were present in Brazil. Species with uncertain nomenclature or questionable records were excluded from the dataset. In addition, the data were supplemented with field collections from the Caatinga region, as well as data from the Cryptogam Herbarium at the State University of Piauí; since this herbarium is not indexed on any online platform, we used a portion of the herbarium’s data. Regarding sample collection, the materials were collected manually using equipment such as a pocket knife, handheld magnifying glass, paper bags and plastic bags, pens, a field notebook, an altimeter, and a GPS device to record geographic coordinates. Samples were stored in kraft paper bags. For macro- and microscopic analysis, we examined the morpho-anatomical characteristics of the gametophyte and sporophyte using a stereoscopic microscope and an optical microscope. For sample handling, we used forceps, needles, scalpels, Petri dishes, slides, and cover slips. We organized all the data into a single database containing information on occurrence, biomes, regions, and Brazilian states where these samples were collected.

#### 4. Project personnel

The principal investigators are Jailton Venilson Ferreira da Silva and Mércia Patrícia Pereira Silva. The associate researchers are Jhonyd Jhonata de Oliveira Marmo, Kleberson Jonata da Cruz Tavares, and Hermeson Cassiano de Oliveira.

## Class III. Data set status and accessibility

### A. Status

1. Latest update: June 26
2. Latest archive date: June 26
3. Metadata status: Last updated on June 26, submitted version.
4. Data verification: Geographic coordinates were verified using Google Earth, and corrections were applied when necessary. Specimens not identified by bryophyte specialists were excluded from the dataset.

### B. Accessibility

1. Contact persons: Jailton Venilson Ferreira da Silva and Mércia Patrícia Pereira Silva. Programa de Pós-Graduação em Biologia Vegetal (PPGBV), Laboratório de Biologia de Briófitas, Departamento de Botânica, Centro de Biociências, Universidade Federal de Pernambuco, Av. Prof. Moraes Rego s/n - Cidade Universitária, Recife, PE 50670-901, Brasil. +55 86 981254170
2. Storage location and medium: The original dataset will be available on the Ecology Repository. The updated version and additional information can be accessed on the Zenodo repository via the following link:
3. Copyright restrictions: Não
4. Proprietary restrictions: Please cite this data paper when using this dataset. Researchers and educators are encouraged to report and share outcomes derived from studies based on these data.
5. Costs: None

## Class IV. Data structural descriptors

A. Data set file

1. Identity: BRYOPHYTES_BRAZIL_Occurrence.csv
2. Size: BRYOPHYTES_BRAZIL_Occurrence.csv, 26 MB
3. Format and storage mode: The dataset is provided as a comma-separated values (CSV) file encoded in UTF-8. Text variables are stored as character strings, whereas geographic coordinates are stored as floating-point numeric values and temporal variables (day, month, and year) are stored as integers. Geographic coordinates are reported in decimal degrees using the WGS84 datum. Missing values are represented as “NA”. Data cleaning procedures included the removal of duplicated records, standardization of taxonomic names, verification of geographic coordinates, and validation of metadata fields.
4. Header information: See column descriptions in section B.
5. Alphanumeric attributes: Mixed
6. Data Anomalies: If no information is available for a given record, this is indicated as ’NA’.
B. Variable information (Table 1).
C. Data anomalies: If no information is available for a given record, this is indicated as ’NA’.

## Class V. Supplemental descriptors

A. Data acquisition
  1. Data acquisition: Our data were obtained from secondary databases and field collections.
  2. Location of completed data forms: No physical forms were used for data collection. The dataset was compiled exclusively from field collections and digital records available in biodiversity information systems and biological collections.
  3. Data entry verification procedures: Data verification involved both automated and manual procedures. Scientific names were standardized according to the adopted taxonomic classification. Geographic coordinates were checked for format consistency and for plausibility of occurrence within Brazilian territory. Duplicate records were identified and removed. Records containing incomplete, inconsistent, or incompatible information relative to the established standards were either corrected when possible or excluded from the final version of the database.
B. Quality assurance/quality control procedures: Several quality control procedures were applied to ensure the taxonomic, geographic, and temporal consistency of the dataset. Scientific names were reviewed and standardized, with synonymy updates and spelling corrections performed when necessary, using widely accepted taxonomic references for bryophytes. Records without species-level identification were removed from the database. Geographic coordinates were checked for decimal degree format (WGS84), and records with invalid coordinates were excluded. The final dataset underwent multiple verification steps to ensure consistency among taxonomic, spatial, and temporal information prior to public release.
C. Related materials: The records used in this study are associated with specimens deposited in national and international herbaria, with relevant information maintained by the respective institutions. Supplementary data include geographic coordinates, taxonomic information, and collection dates. The dataset does not include photographs, videos, or original field notebooks.
D. Archiving
  1. Archival procedures: The final version of the dataset will be made available in CSV format in a public open-access repository, ensuring long-term digital preservation, traceability, and permanent access to the data used in this study.
  2. Redundant archival sites: In addition to the version deposited in the primary repository, copies of the dataset are maintained by the authors.
E. Publications and results: The dataset was compiled to support studies on the diversity, geographic distribution, and macroecological patterns of bryophytes in Brazil. The associated results will be presented in this Data Paper and may support future research in biodiversity, biogeography, conservation, and bryophyte ecology. Publications derived from this dataset should appropriately cite this Data Paper and the repository where the data are archived.
F. History of data set usage
  1. Data request history: None
  2. Data set update history: None
  3. Review history: None
  4. Questions and comments from secondary users: None

## ACKNOWLEDGMENTS

We thank Coordenação de Aperfeiçoamento de Pessoal de Nível Superior (CAPES; Finance Code 001) and Conselho Nacional de Desenvolvimento Científico e Tecnológico (CNPq; Universal Grant No. 409503/2025-8 and PPBio-RABECA Network Grant No. 441209/2023-8).

---

**Table 1.** Description of the fields contained in the file Brazilian_Bryophytes_Occurrences.csv. The dataset comprises 102,527 occurrence records of bryophytes from Brazil compiled from multiple biodiversity databases, herbarium collections, and published sources.

| Variable | Description | Levels/Units | Example |
|---|---|---|---|
| Source | Source of occurrence record | 4 data sources | GBIF |
| Record_Id | Record identification in each data set. Authors were free to use their own identification | The data set comprises 102,527 record IDs. | BRYO_000001 |
| Phylum | Bryophyte phylum | 3 phyla | Bryophyta |
| Family | Taxonomic family | 131 families | Fissidentaceae |
| Species | Accepted scientific name | 1,835 species | Fissidens crispus |
| Locality | Locality description provided by the source | 23,272 localities | Recife |
| Latitude | Geographic latitude (WGS84) | Decimal degrees | -8.0522 |
| Longitude | Geographic longitude (WGS84) | Decimal degrees | -34.9286 |
| day | Day of collection | 1–31 | 15 |
| month | Month of collection | 1–12 | 7 |
| Year | Year of collection | 184 unique years | 2018 |
| catalogNumber | Catalog number assigned by the collection | 90,318 catalog numbers | PEUFR12345 |
| recordNumber | Collector record number | 30,151 record numbers | 234 |
| identifiedBy | Person responsible for taxonomic identification | 3,170 identifiers | O. Yano |
| recordedBy | Collector(s) of the specimen | 5,867 collectors | J. P. Frahm |
| institutionCode | Institution acronym | 178 institutions | PEUFR |
| collectionCode | Collection acronym | 154 collections | Bryophytes |
| rightsHolder | Institution or person holding data rights | 136 rights holders | Universidade Federal de Pernambuco |
| state | Brazilian state of occurrence | 27 states | Pernambuco |
| region | Brazilian geographic region | 5 regions | Northeast |
| biome | Brazilian biome | 7 biomes | Atlantic Forest |

